# Application of adaptive-network-based fuzzy inference systems to the parameter optimization of a biochemical rule-based model

**DOI:** 10.1101/338384

**Authors:** Brittany R. Hoard

## Abstract

Our main contribution is an efficient machine learning approach to fitting parameters of a biological model. We study the binding of the shrimp protein Pen a 1 with antibody-receptor complexes because this process is important in understanding the allergic response. Previously, we developed a BioNetGen model that simulates this process. We previously developed a method for encoding steric effects via the optimization of two parameters: the cutoff distance and the rule rate. We optimized these two parameters by fitting the output to that generated by a 3D robotics-inspired Monte Carlo simulation that explicitly represents molecular geometry.

In this work, we aim to optimize the parameters for our BioNetGen model using an efficient method: an adaptive-network-based fuzzy inference system implemented in MATLAB. We want to develop fuzzy systems that can accurately predict the rule binding rate and cutoff distance given a residual-sum-of-squares value or a probability distribution. We construct the fuzzy systems using fuzzy c-means clustering with existing data from BioNetGen model parameter scans as the training data. We create and test fuzzy systems with various input data and number of clusters, and analyze their performance with regard to the effective optimization of our rule-based model. We find that the fuzzy system that uses a residual-sum-of-squares value as the input value performs acceptably well. However, the performance of the fuzzy systems that use probabilities as their input values perform inconsistently in our tests.

The results of this study suggest that the system that uses a residual-sum-of-squares value as the input value could potentially be used to find an adequate fit for our biochemical model. However, the systems that use probabilities as their input values need further development to improve the consistency and reliability of their output. Testing more values for other clustering parameters other than the number of clusters may accomplish this. Further research could also include similar studies using other training or clustering algorithms. This methodology could be modified for use with fitting other biological models.

## 1 BACKGROUND

### 1.1 Introduction

Fuzzy inference systems (FISs) are models that consist of a set of IF-THEN rules in which the antecedents and/or consequents of the rules are fuzzy rather than crisp Wang and Mendel (1992). The rules can be provided by a human expert, but it can be highly useful for the rules to be constructed automatically using only a set of training data and an appropriate learning algorithm. Various FIS learning algorithms have been developed. FISs can be applied to regression and classification problems in various fields including robotics, data mining, prediction, estimation, control, and computational biology.

In this paper, we examine a novel application of FISs in the field of biology; namely, the optimization of the parameters of a biochemical rule-based model implemented in BioNetGen Blinov et al. (2004) which could potentially be modified for implementation in other biological models. Ideally, this method would require only one parameter scan to generate the set of data used to train the FIS. In general, BioNetGen models consist of a set of “rules" that control interactions between molecules. In a previous line of work, we developed a BioNetGen model that captures the binding interactions between molecules of the shrimp tropomyosin Pen a 1 and IgE antibody-receptor complexes that lead to the formation of aggregates. We are interested in this biological process because the size and structure of these aggregates is important in understanding the allergic response in shrimp-allergic human subjects as aggregate size and structure is theorized to be linked to the strength of the allergic response. Such knowledge could be useful in developing treatments for people with allergies, such as the administration of recombinant hypoallergens Vargas et al. (2018). Using this BioNetGen model, we want to determine the probability that an aggregate of a particular size will be formed. We use the term *aggregate size* to refer to the number of IgE antibody-receptor complexes bound to a single Pen a 1 molecule. We use the BioNetGen model output to calculate these probabilities.

The Pen a 1 allergen is a shrimp tropomyosin. It is a dimer; it has a double-stranded coiled structure. In previous experimental work, ten binding regions of the double-stranded Pen a 1 molecule have been identified (five per strand) Ivanciuc et al. (2003); Ayuso et al. (2002); Reese et al. (2005). However, in previous work performed by colleagues, one large binding region was effectively split into two regions based on a study of conditional binding probabilities of these two regions Manavi et al. (2016). Consequently, we treat each Pen a 1 molecule as having a total of 12 binding regions (six per strand). This means that there are 13 possible aggregate sizes (0-12 regions can be bound).

We wish to optimize two parameters of our biochemical rule-based model: the cutoff distance and the rule binding rate, which we refer to throughout this paper as the “rule rate". The *cutoff distance* is the distance between tropomyosin binding sites at or below which steric effects become strong enough to significantly reduce the probability of a binding event taking place at a neighboring tropomyosin binding site. We encode steric effects between the binding sites by changing the set of rules according to the cutoff distance. The *rule rate* is the probability that an event encoded by that rule will occur. For our simple model, we make the assumption that the same rate is associated with each rule. Previously, the determination of the rule rates for the rule-based model has been achieved by parameter scanning, which can be time-consuming and risks skipping over the best fit if the step size is too large. An algorithm that uses the Metropolis method has also been used, though it can also be time-consuming. FISs only take a maximum of a few minutes to train for our model, and they can then produce output instantaneously given a set of input values. The greater efficiency and possibly greater accuracy of the FIS method of parameter optimization are the motivations for trying to apply this method to the optimization of our BioNetGen model.

A common reason to optimize the parameters of a biological model is to fit the model output to a set of experimental data. Because experimental data for this particular biological process is not currently available, we substitute aggregate size data from a three-dimensional rigid-body Monte Carlo simulation previously developed by our collaborators Manavi et al. (2012). We seek to optimize the rule rate and cutoff distance for our biological model to reduce the difference between the aggregate size distribution of the Monte Carlo data and that of our rule-based model. We quantify this difference by calculating the residual sum-of-squares (RSS) value between the aggregate size distribution generated by our model and that generated by the Monte Carlo simulation.

There are multiple options for designing an FIS for this optimization problem since there are 13 possible aggregate sizes, a cutoff distance, a rule rate, and an RSS value associated with each training data point. All of these parameters could be used as input parameters to the FIS. We could potentially construct a single-input FIS with only one input parameter, or a multiple-input FIS with more than one input parameter. The main contributions of this work are the creation and testing of different FISs, including single-input and multiple-input systems, and an analysis of their performance with regards to the effective optimization of our rule-based biological model for allergen-antibody aggregation implemented using BioNetGen. We came to the conclusion that the single-input RSS system performs acceptably well, but the performance of the multiple-input systems is inconsistent, with one system performing well and the other systems performing poorly.

### 1.2 Related Work

There have been numerous learning algorithms developed for FISs. One of the earliest and most widely used algorithms is the Wang-Mendel technique Wang and Mendel (1992), in which the input and output spaces are divided into fuzzy regions that form the basis of the fuzzy rules, and each rule is assigned a degree of usefulness. The adaptive-network-based fuzzy inference system (ANFIS) Jang (1993); Jang et al. (1997) is a two-stage model. The forward stage consists of multiple layers, including fuzzification, inference, and other calculations, and parameter learning using the least-squares method takes place in the backward stage. In the subtractive clustering and fuzzy c-means method Chiu (1996); Yager and Filev(1994), rule cluster centers are determined by calculating the distance of each data point from every other data point, and then optimizing the clusters using fuzzy c-means. The MOGUL method Herrera et al. (1998); Cordon et al. (1999) uses iterative rule learning to generate chromosomes that consist of the rule membership function parameters. In the fuzzy inference rules by descent method Nomura et al. (1992), the antecedent membership function is an isosceles triangle, and the consequent part of the rule is a real number obtained using a descent method.

### 1.3 Contributions

The contributions of this work are as follows:

1. The proposal of an efficient fitting method for biological models using fuzzy inference systems.
2. The application of this method to the problem of fitting the parameters of a biological rule-based model.
3. The testing of the accuracy of this method for finding reasonable fits and consistency of the results.

## 2 METHODS

For our FIS learning algorithm, we selected the adaptive-network-based fuzzy inference system (ANFIS) for its implementation and customization options in MATLAB. An initial fuzzy inference system (FIS), which includes the fuzzy rule base and membership functions, is generated using fuzzy c-means clustering. The parameters of this initial system are then trained further using ANFIS. Further detail regarding the construction of the FIS can be found in the online MathWorks documentation MathWorks (2018). One parameter of particular interest is the number of clusters used in fuzzy c-means clustering. During this process, clusters are identified within the training data, and these clusters are then employed in the generation of the FIS. The number of clusters may be specified by the user and can have a significant effect on the results, so this study includes systems with various numbers of clusters used.

Ideally, we want to develop two FISs: one that can accurately predict the rule rate, and another that can accurately predict the cutoff distance, given an aggregate size probability distribution or an RSS value as the input value. Having such FISs would be helpful for optimization of our BioNetGen model, as ideally, only a small set of training data would be needed to train the FISs. These FISs could then be fed in the Monte Carlo aggregate size probability distribution as the input values and would accurately predict the rule rate and cutoff distance that best corresponds to that particular distribution. This method would be far less time-consuming than multiple-parameter scans and even Metropolis-based algorithms.

For the application of FISs to our BioNetGen model, the ANFIS method was chosen based on its easy-to-use implementation in MATLAB and its customization options. This method employs an adaptive network that is composed of nodes and directional links that connect the nodes, and at least some of the nodes are adaptive (the nodes are associated with parameters that are adapted to minimize an error of measure according to the learning rules of the network) Jang (1993). ANFIS uses hybrid learning rules that combine the gradient method and the least squares estimate Jang et al. (1997).

The application of FISs to the BioNetGen model is based on previous work using a 3D geometric Monte Carlo simulation to model the aggregation of IgE antibodies onto the shrimp allergen tropomyosin Manavi et al. (2016). For this simulation, isosurface models of Pen a 1 were created from all-atom structures of shrimp tropomyosin. These structures were obtained from the Protein Data Bank (PDB:1CG1) and the Structural Database of Allergenic Proteins (SDAP Model #284) Ivanciuc et al. (2003, 2002). The tropomyosin molecule used contains 568 amino acids and 4,577 atoms. It is a dimer and has a double-stranded coiled structure.

A rule-based model implemented in BioNetGen that implicitly represents antigen geometry was used to quantify the differences in the Monte Carlo results for different allergen conformations and resolutions of the Monte Carlo model. This work employs a method to construct sets of rules based directly on the distances between the IgE binding regions of the tropomyosin Hoard et al. (2015,2016). Once the cutoff distance is specified, the set of rules is constructed according to the cutoff distance and the distances between the binding regions. Hence, the number of rules in the model varies according to the cutoff distance.

We use the ANFIS method, implemented in MATLAB, to train the FISs. The FISs will be constructed using fuzzy c-means clustering with existing aggregate size data from BioNetGen model parameter scans as the training data. Either an RSS value or some of the 13 possible aggregate size probabilities will be used as input variables, and a rule rate or cutoff distance will be the output variable. For the final test, all of the Monte Carlo aggregate size probabilities will be used as input variables to the FIS.

The performance of the FISs will be evaluated by calculating the percent error between the output values generated by the FIS and the actual values, and by simply observing the output values and seeing if they are reasonable for our expectations of the particular model. This point is further discussed in the Results section. For the final test, the FIS will be evaluated by comparing the RSS values to the minimum RSS value predicted by the Metropolis algorithm.

### 2.1 Computational Experiments

In order to construct a fuzzy system using ANFIS, the genfis3 function in MATLAB is first run to create a Sugeno-type FIS structure using fuzzy c-means clustering to extract a set of rules and membership functions that model the training data. This function allows the specification of the number of clusters used to model the data. This parameter will be varied throughout this study. The other parameters were set to their default values; the number of training epochs was set to 10, the initial step size was set to 0.01, the step size increase rate was set to 1.1, and the step size decrease rate was set to 0.9.

The data used to train the FIS consists of the rule rates, the aggregate size probability distributions, the cutoff distances, and the RSS values. This data was obtained by running a parameter scan of the rule rate from 0.000 *molecule*^−1^ *s*^−1^ to 0.020 *molecule*^−1^ *s*^−1^ with a step size of 0.001 *molecule*^−1^ s^−1^ over a range of cutoff distances from 3.5 nm to 10.0 nm with a step size of 0.1 nm. This training set consists of 1,365 data points and is provided in the supporting information for this article.

Before developing the more complex multiple-input FIS, there is another type of FIS we can develop that could be applied to optimization: a single-input FIS that uses an RSS value as an input variable and the rule rate or cutoff distance as the output variable. The disadvantage of this FIS is that we do not know what the minimum possible RSS value is; however, we do have an idea from our experience with our biological models of what a “good" RSS value should be, and we can try using different “good" RSS values to find an accurate rule rate or cutoff distance. The advantage of this FIS is that it is far less computationally expensive than a multiple-input FIS.

In order to determine how well the FIS can predict the rule rate and cutoff distance, we first randomly select five RSS values within the range of the training data (but with none matching that of any of the training values). Since the lowest RSS value is likely to be outside of the range of the training data, we also want an idea of how well the FIS performs with this type of input. Two small RSS values below the lowest RSS value in the range of the training data are chosen for this purpose. After training the FIS, we use each of these seven RSS values as the input variable and use the FIS to predict the rule rate and cutoff distance for each of these seven values. We then run a simulation for our BioNetGen model using the rule rate and cutoff distance predicted by the FIS and note the actual RSS value calculated for the BioNetGen simulation. Ideally, the inputted and predicted RSS values should be similar.

## 3 RESULTS

### 3.1 Single-Input FIS

Figure 1 and Figure 2 show the results of this test. Figure 1 displays the predicted RSS values for the five randomly selected inputted RSS values and the two non-randomly selected inputted RSS values. In this figure, the inputted RSS values are plotted on the x-axis, and the RSS values predicted by the FIS are plotted on the y-axis. There are five curves, one for each of the number of clusters used in the initial construction of the FIS. Figure 2 displays the percent error between the inputted and predicted RSS values for each number of clusters.

**FIGURE 1.**
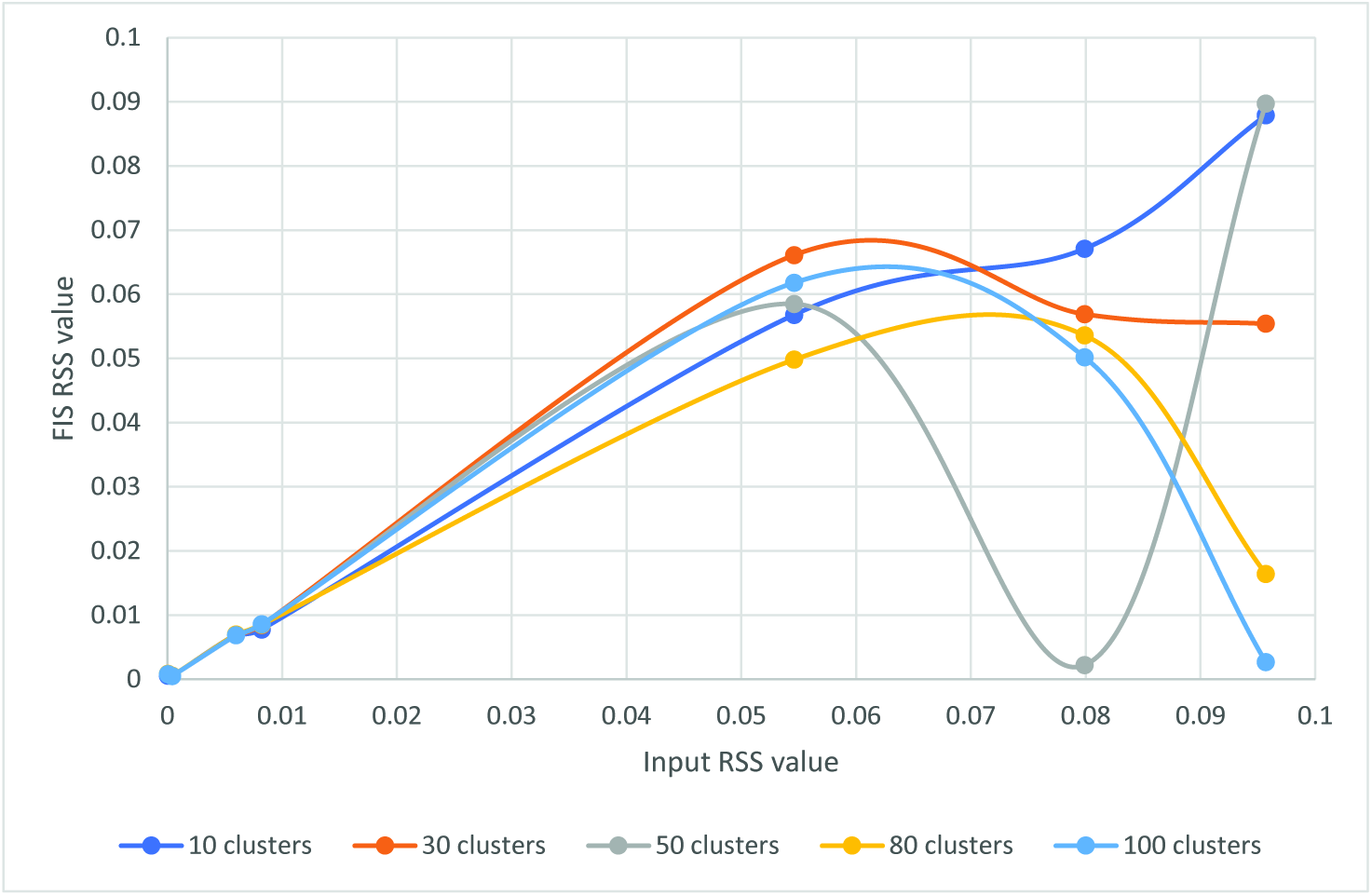
RSS values for the BioNetGen models with the rule rates predicted by the FIS with various numbers of clusters using the selected RSS values as input.

**FIGURE 2.**
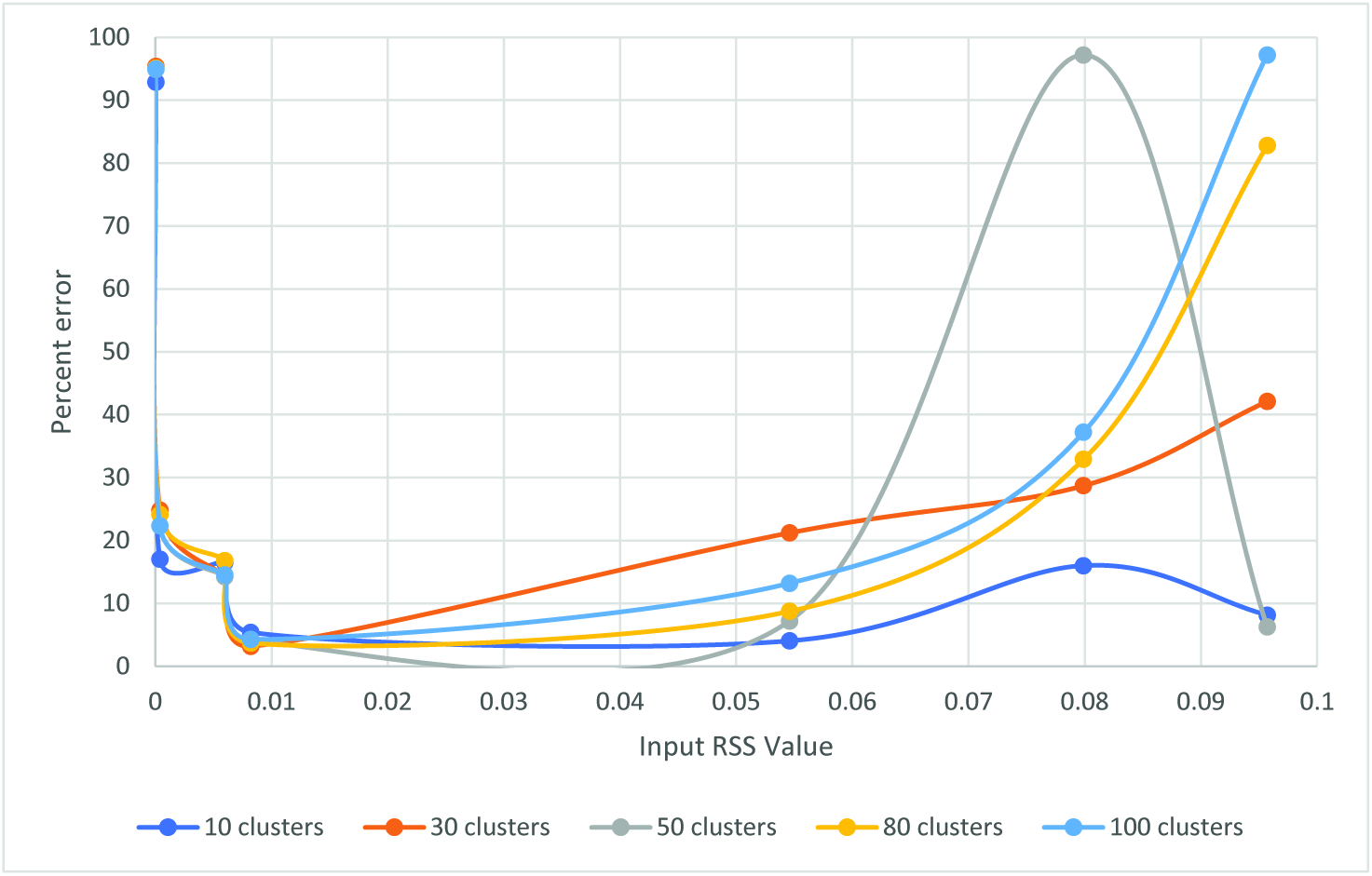
Percent error between the inputted and predicted RSS values.

From Figure 1, we can observe that the FIS performs the best for this test with only 10 clusters. We also observe from Figure 2 that the system performs especially poorly for the higher RSS values, with errors of over 90% for two tests. This could be due to the fact that a high RSS value may correspond to a large range of poorly fitting data sets, while a lower RSS value corresponds with a much smaller number of well-fitting data sets.

The lowest RSS value, 0.00004, also performs poorly, probably because this RSS is too small to be achievable. It is well outside the range of the training data. The system performs reasonably well for the next lowest RSS input value, 0.0004, which is also outside the range of the training data, but close to the minimum value found by the Metropolis algorithm, 0.000481558. Many of the predicted RSS values are close to this Metropolis value, which is an indication that the performance of this FIS is similar to that of our Metropolis fitting algorithm. Since the predicted RSS values for these two input values are all less than 0.001, and since we ultimately seek to use this system to find reasonably good fits (generally corresponding to RSS values less than 0.001), this performance is acceptable.

#### Random Rule Rates

The purpose of this test is to determine how accurately the FIS can predict a rule rate given a set of aggregate size probabilities corresponding to that rule rate. Firstly, five rule rates were selected from a uniform distribution in the interval [0.00,0.02] *molecule*^−1^*s*^−1^, and were each specified as the variable rule rate for the same BioNetGen model.

The rule base varies with the cutoff distance, so for this initial test, a cutoff distance of 4.7 nm was specified for all five BioNetGen models. This distance was chosen because our Metropolis fitting algorithm found that the best fitting BioNetGen model has this cutoff distance. The BioNetGen aggregate size probability distributions were generated for each of the five rule rates. The probability values for aggregate sizes 5 through 10 were then fed into the FIS, which was used to predict the rule rates. (Not all 13 of the size values were used as input variables because 13 input variables makes the model computationally intractable. Sizes 5 though 10 were selected because these sizes are most likely to correspond to values that are large enough to be significant.) These predicted rule rates should ideally be similar to the actual rule rates used to generate the BioNetGen model.

Table 1 and Figure 3 show the results of this test.

**TABLE 1.** Rule rates predicted by the FIS compared with the actual rule rates

| Selected rate | Clusters |  |  |  |  |
| --- | --- | --- | --- | --- | --- |
|  | 10 | 30 | 50 | 80 | 100 |
| $3.26 \times 10^{-3}$ | $3.01 \times 10^{-3}$ | $3.26 \times 10^{-3}$ | $3.26 \times 10^{-3}$ | $3.27 \times 10^{-3}$ | $3.29 \times 10^{-3}$ |
| $7.43 \times 10^{-3}$ | $7.02 \times 10^{-3}$ | $7.47 \times 10^{-3}$ | $7.42 \times 10^{-3}$ | $7.42 \times 10^{-3}$ | $7.43 \times 10^{-3}$ |
| $7.91 \times 10^{-3}$ | $7.43 \times 10^{-3}$ | $8.07 \times 10^{-3}$ | $7.92 \times 10^{-3}$ | $7.91 \times 10^{-3}$ | $7.91 \times 10^{-3}$ |
| $9.76 \times 10^{-3}$ | $9.88 \times 10^{-3}$ | $9.69 \times 10^{-3}$ | $9.75 \times 10^{-3}$ | $9.76 \times 10^{-3}$ | $9.77 \times 10^{-3}$ |
| $1.78 \times 10^{-2}$ | $1.78 \times 10^{-2}$ | $1.78 \times 10^{-2}$ | $1.78 \times 10^{-2}$ | $1.78 \times 10^{-2}$ | $1.78 \times 10^{-2}$ |

**FIGURE 3.**
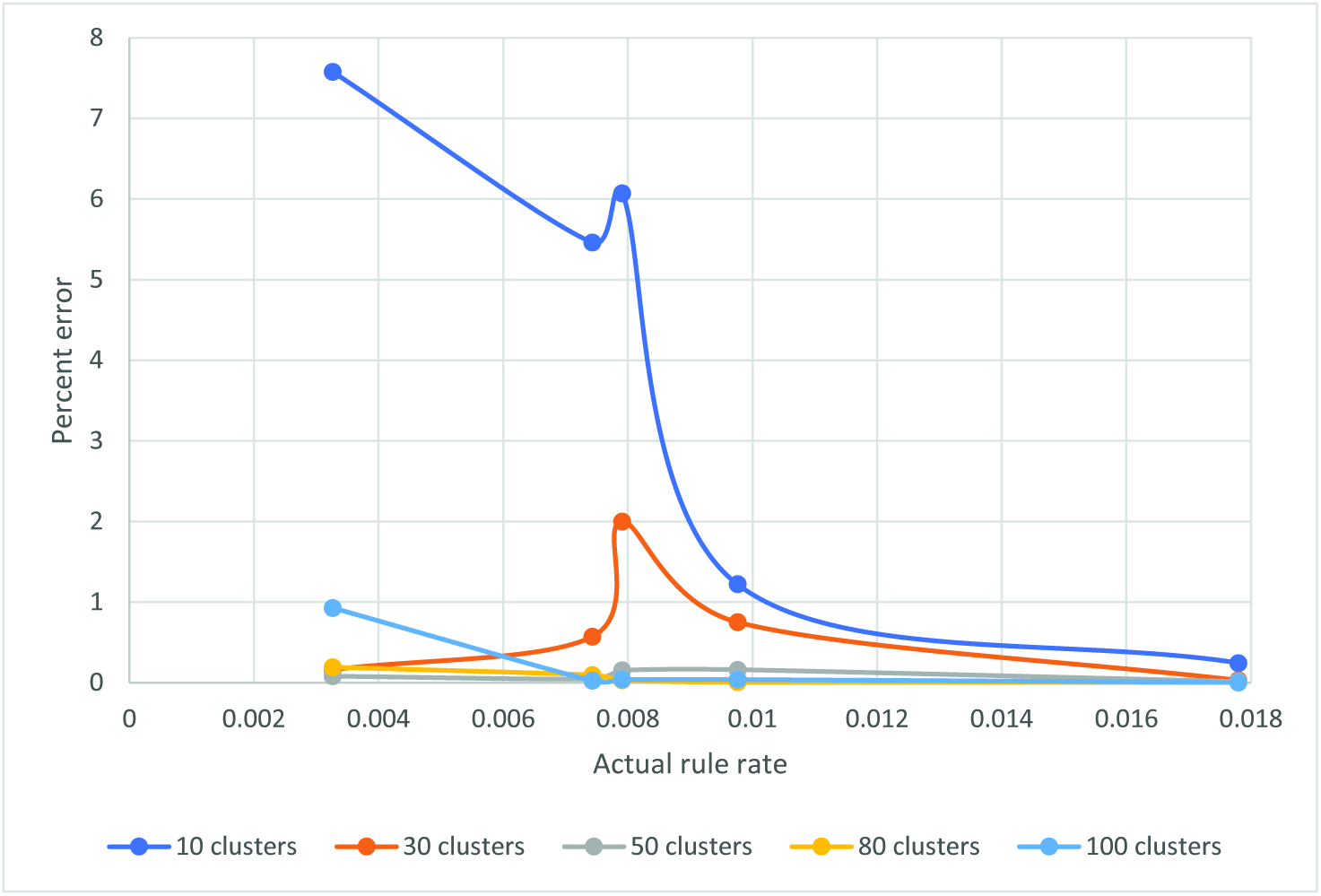
Percent error between the actual and predicted rule rates.

Table 1 and Figure 3 show that this FIS performs well at predicting the rule rates given a set of aggregate size probabilities generated by BioNetGen, with most of the percent error values being less than one. We also observe that the system performs better with a higher number of clusters for this test.

### 3.2 Random Cutoff Distances

The purpose of this test is to determine how accurately the FIS can predict a cutoff distance given a set of aggregate size probabilities corresponding to that cutoff distance. Firstly, five cutoff distances were selected from a uniform distribution in the interval [3.5,10.0] nm, and were each specified as the cutoff distance for a BioNetGen model with the same rule rates. For this test, the variable rule rate was specified as 0.00983 *molecule*^−1^ *s*^−1^ for all models. This rule rate was chosen because our Metropolis fitting algorithm found that the best fitting BioNetGen model has this rate. The aggregate size probability distributions were generated for each of the five cutoff distances. These probability values were then fed into the FIS, which predicted the cutoff distances. These predicted cutoff distances should ideally be similar to the actual cutoff distances used to generate the BioNetGen model.

Table 2 and Figure 4 show the results of this test.

**TABLE 2.**
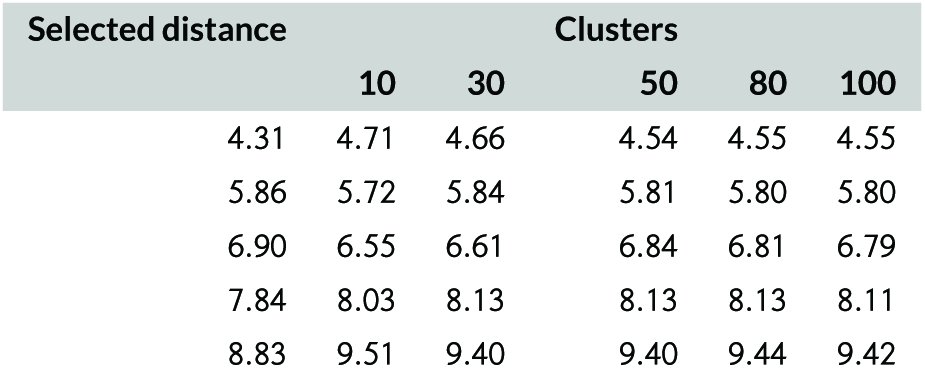
Cutoff distances predicted by the FIS compared with the actual cutoff distances.

| Selected distance | Clusters |  |  |  |  |
| --- | --- | --- | --- | --- | --- |
|  | 10 | 30 | 50 | 80 | 100 |
| 4.31 | 4.71 | 4.66 | 4.54 | 4.55 | 4.55 |
| 5.86 | 5.72 | 5.84 | 5.81 | 5.80 | 5.80 |
| 6.90 | 6.55 | 6.61 | 6.84 | 6.81 | 6.79 |
| 7.84 | 8.03 | 8.13 | 8.13 | 8.13 | 8.11 |
| 8.83 | 9.51 | 9.40 | 9.40 | 9.44 | 9.42 |

**FIGURE 4.**
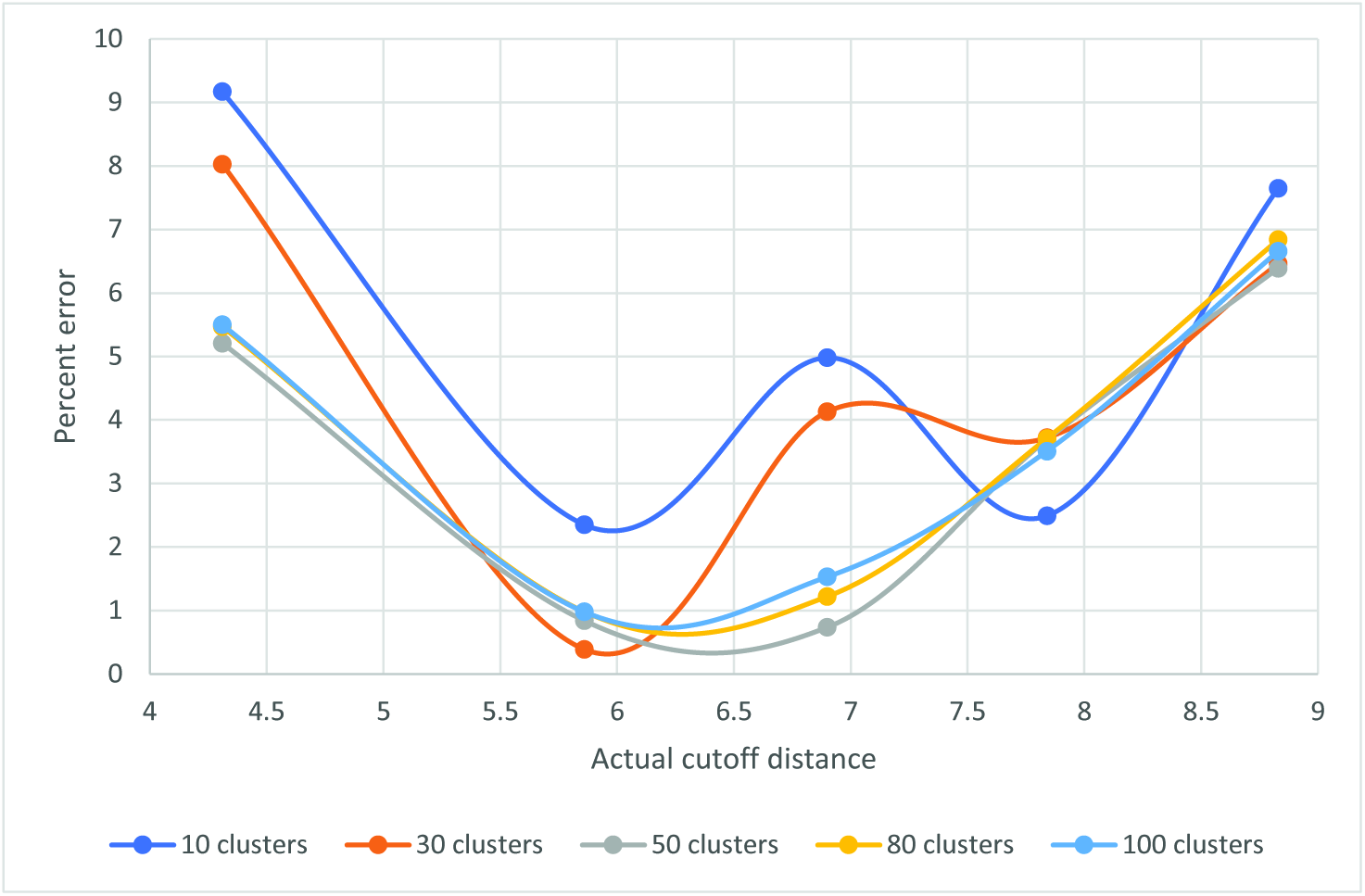
Percent error between the actual and predicted cutoff distances.

Table 2 and Figure 4 show that this FIS performs reasonably well at predicting cutoff distances given a set of aggregate size probabilities generated by BioNetGen, although some of the error values are rather high, with a few greater than five percent and one greater than nine percent. This could potentially pose a problem for this system, since we seek a cutoff distance accurate to within 0.1 nm according to the cutoff distance step size. As it stands, this FIS is better suited to finding an approximate “best" cutoff distance that can then be used to select a small range of cutoff distances that can be further optimized to find the true best cutoff distance (and rule rate). We also observe that the system tends to perform better with a higher number of clusters for this test.

### 3.3 Monte Carlo Model Prediction

The final test of our FIS is whether the Monte Carlo data can be used as input variables to predict a rule rate and cutoff distance that correspond to a good fit. In this test, a set of Monte Carlo aggregate size probabilities is fed into the FIS, and the rule rate and cutoff distance are predicted. The predicted rule rate and cutoff distance are tested by using these values as the rule rate and cutoff distance in the BioNetGen model, generating the aggregate size distribution from BioNetGen, and comparing this distribution to the Monte Carlo data. Since we do not know what the optimal RSS value is, we compare the results to that of the Metropolis-based optimization algorithm to see how well the FIS compares.

Table 3 shows the results of this test.

**TABLE 3.**
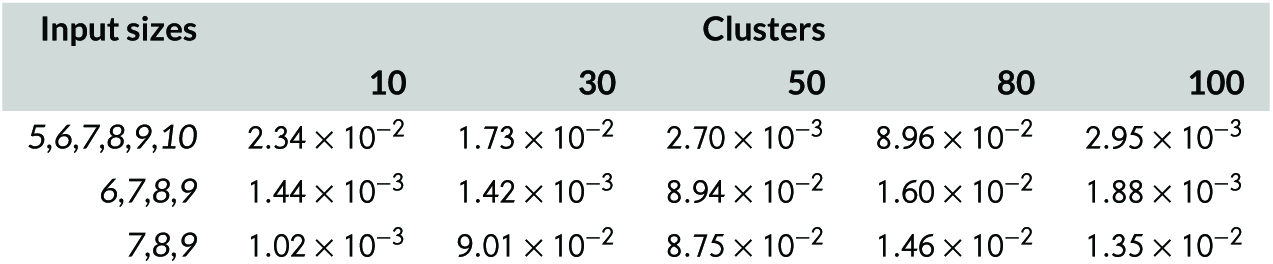
RSS values for the BioNetGen model using the FIS-predicted rule rate and cutoff distance. The leftmost column represents the Monte Carlo aggregate sizes used as input variables.

We note that the minimum RSS value found for this BioNetGen model using a Metropolis-based algorithm is 0.000481558. Comparing this value with the results in Table 3, we observe that all of the FIS-predicted values are at least one order of magnitude higher than this value. We also note that the performance of this system is inconsistent, and many RSS values are unacceptably high (greater than 0.01). This FIS needs improvement before it can be used as a tool for finding best fits.

## 4 CONCLUSIONS

In this study, we sought to create an FIS that can accurately predict best-fit rule rates and cutoff distances for a BioNetGen rule-based model. We tested different FISs using (a) an RSS value, (b) a set of aggregate size probabilities, and (c) a set of aggregate size probabilities directly from the Monte Carlo simulation data as the FIS input variables. A FIS corresponding to (a) or (c) would be especially useful for this optimization problem. Our FIS based on (a) performed well for low RSS values and could potentially be used to predict rule rates and cutoff distances that result in a good fit for the BioNetGen model (although it will not necessarily find the best possible fit, it could come very close). The FISs based on (b) consistently performed well at predicting rule rates, but its prediction of cutoff distances was rather inaccurate. However, this system could still be used to narrow down the range of cutoff distances. Finally, the FIS based on (c) performed poorly overall. We also note that the optimal number of clusters varies depending on the type of input. For most tests, a higher number of clusters was linked to better performance of the FIS, although this was not always the case.

Our results suggest that the use of an FIS for fitting parameters of a biological model has the potential to be effective and efficient. Future work on this problem could involve testing other fuzzy training algorithms besides ANFIS and other clustering or rule construction algorithms besides fuzzy c-means clustering. Other ANFIS and clustering parameters besides the number of clusters, such as change in step size rate, training epoch number, and initial step size, could be tested to see if any of these parameters have a significant effect on the FIS performance.

## Abbreviations

ANFIS: Adaptive network based fuzzy inference system
FIS: Fuzzy inference system
nm: Nanometers
RSS: Residual sum of squares
s: Seconds

## ACKNOWLEDGEMENTS

Thanks to the University of New Mexico Department of Computer Science for providing the funding that made this work possible. Thanks to the faculty and fellow students who asked insightful questions about this work and contributed to the improvement of the manuscript.

